# UFCG: database of universal fungal core genes and pipeline for genome-wide phylogenetic analysis of fungi

**DOI:** 10.1101/2022.08.16.504087

**Authors:** Dongwook Kim, Cameron L.M. Gilchrist, Jongsik Chun, Martin Steinegger

## Abstract

In phylogenomics the evolutionary relationship of organisms is studied by their genomic information. A common approach to phylogenomics is to extract related genes from each organism, build a multiple sequence alignment and then reconstruct evolution relations through a phylogenetic tree. Often a set of highly conserved genes occurring in single-copy, called core genes, are used for this analysis, as they allow efficient automation within a taxonomic clade. Here we introduce the Universal Fungal Core Genes (UFCG) database and pipeline for genome-wide phylogenetic analysis of fungi. The UFCG database consists of 61 curated fungal marker genes, including a novel set of 41 computationally derived core genes and 20 canonical genes derived from literature, as well as marker gene sequences extracted from publicly available fungal genomes. Furthermore, we provide an easy-to-use, fully automated and open-source pipeline for marker gene extraction, training and phylogenetic tree reconstruction. The UFCG pipeline can identify marker genes from genomic, proteomic and transcriptomic data, while producing phylogenies consistent with those previously reported, and is publicly available together with the UFCG database at https://ufcg.steineggerlab.com.

## Introduction

The taxonomic kingdom *Fungi* is one of the most diverse clades in the tree of life, potentially encompassing 2.2 to 3.8 million species (1). Publicly listed fungal species in the Ref-Seq database (2) have exponentially grown from only 157 to 16,869 species in the last 12 years. Fungal resources, such as available genomes or proteomes are growing rapidly and are powering the resolution of phylogenetic relationships within the clade.

Before this wealth of fungal resources became available, only few near universally present markers were used for phylogenetic analysis. The internal transcribed spacer (ITS) region of the nuclear ribosomal RNA (rRNA) cistron has long been used in phylogenetic analysis of fungi as the universal fungal marker (3–5) and has formed the basis of large-scale barcoding efforts (6, 7). In cases where the ITS region does not provide adequate resolution, secondary markers may be used instead (8), such as RNA polymerases (9), translation elongation factors (10), and mitochondrial genes (11). The use of multi-gene phylogenies, where the ITS region is used in conjunction with these secondary markers, has become increasingly common to resolve taxonomic relationships (12–15).

However, usage of secondary markers requires researchers to know which markers to select based on the lineage being studied, as well as how to practically extract, align, and concatenate them to infer their phylogenetic relationship (16).

With the rapid increase of available public genomic sequences analysis using multiple markers became feasible, which increased the resolution further (17). The most commonly used technique is to concatenate multiple genes that are single-copy and orthologous, while existing universally among the taxa (core genes; 18). Core gene based automated phylogenomic pipelines have attained wide adoption in the prokaryotic kingdom, such as Genome Taxonomy Database (GTDB; 19), AutoMLST (20), or UBCG (21).

There have been numerous efforts towards defining sets of single-copy orthologs for the fungal kingdom, notably the Fungal Genome Mapping Project (FGMP; 22), and the Benchmarking Universal Single-Copy Orthologs pipeline (BUSCO; 23), which implements OrthoDB datasets (24). However, the primary focus of these methods is on assessing the completeness of fungal genomes, and as such no single method integrates the entire process from core gene identification to phylogeny reconstruction.

Here, we introduce the Universal Fungal Core Genes (UFCG), a database of fungal marker genes derived from experimentally annotated genes (Fig. 1a) as well as a pipeline for genome-wide phylogenetic analysis (Fig. 1b). We defined 61 marker genes, 20 canonical markers extracted from literature research and 41 marker genes determined by computational identification of single-copy and highly conserved genes across the fungal tree of life, starting from manually curated and well-annotated sequences. The UFCG database provides freely accessible resources about the marker genes and fungal species from which the markers were identified, accompanied by a user-friendly website. We also provide an easy-to-use and fully integrated pipeline for fungal marker gene profiling and phylogenetics from fungal genomic, transcriptomic, and proteomic data.

**Fig. 1.**
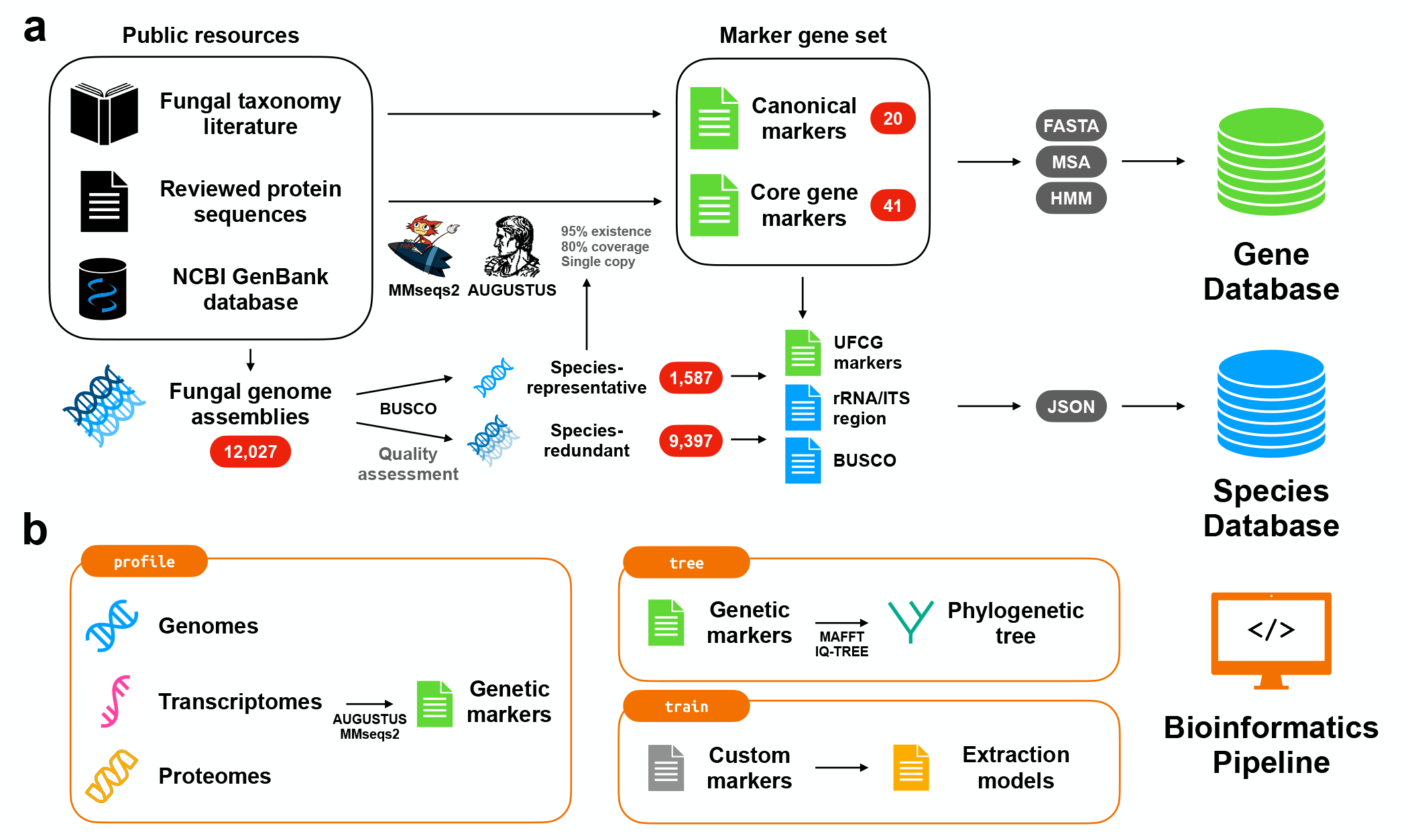
Schematic illustration of the preparation of the UFCG database and pipeline. (a) The UFCG Gene database consists of novel 41 core gene markers we defined, and 20 canonical marker genes curated from fungal taxonomy literature. We built profiles for all SwissProt Fungi proteins and searched them against 1,587 species-representative genome assemblies using MMseqs2. Only genes that occur as single-copy in at least 95% species were further refined and filtered by AUGUSTUS-PPX. For each gene, we offer profile hidden Markov models (HMMs) and the seed amino acid sequences, downloadable from the database. The UFCG Species database provides pre-extracted marker sequences from the genome assemblies we obtained. In addition to the marker genes we defined, we extracted ITS and BUSCO sequences from both 1,587 species-representative and 9,397 species-redundant fungal genome assemblies. We compiled the extracted sequences into JSON files, which are downloadable from the database. (b) Graphical representation of three main modules (`profile`, `tree`, and `train`) from the pipeline. The `profile` module accepts genomic, proteomic, and transcriptomic data of fungi and extracts marker sequences using a pre-trained set of profile HMMs. The `tree` module combines the set of extracted marker genes and reconstructs their phylogeny as a maximum likelihood tree using aligned and concatenated marker sequences. The `train` module converts custom marker sequences into profile HMMs that can be directly utilised by the `profile` module.

## Materials and Methods

### Preparation of genome assemblies

We obtained 12,027 whole genome assemblies of fungal species from the National Center of Biotechnology Information (NCBI) GenBank database (25) (Fig. 1a). Taxonomic information from the NCBI taxonomy database (26) was assigned to each assembly. A single genome assembly was chosen for each fungal species by selecting those marked as representative in the GenBank database, excluding species with non-unique nomenclature (e.g., fungal *sp*.).

The completeness of the remaining assemblies was assessed by searching for single-copy orthologs using BUSCO v3.0.2 (27) with the OrthoDB v9 fungal lineage dataset (28). Additionally, we performed *ab initio* gene prediction using AUGUSTUS v3.4.0 (29) with a pre-trained species model of *Rhizopus oryzae*, which resulted in the highest average prediction count among the models (Supplementary Table 1). Assemblies that failed to report ≥ 250 BUSCO and ≥ 3,000 predicted genes were removed from the set, resulting in 1,587 species-representative assemblies.

### Core gene candidate detection with MMseqs2

For our core gene candidates we started with accurately annotated and experimentally validated genes from Swiss-Prot to avoid incorrectly called genes from contaminated genomic fragments (30) or fragmented genes because of limitations of eukaryotic gene finding software (31).

All 35,591 fungal protein sequences present in Swiss-Prot (release 2022_03; 32), the manually curated part of the UniProt KnowledgeBase (33), were extracted and clustered to 90% sequence identity using MMseqs2 v13.45111 (34), resulting in 30,834 representative sequences.

For each sequence, we generated a query centered multiple sequence alignment (MSA) by searching for three iterations (`--num-iterations 3`) against the full UniProtKB release 2022_03 (35) using MMseqs2. Each MSA was turned into a profile and searched against the species-representative assemblies using a MMseqs2 six-frame-translated sequence-to-profile search.

In some cases, hits to certain genes may be fragmented into multiple smaller hits (e.g., due to the intron-exon structure of eukaryotic genes), causing them to be filtered out in downstream analyses. In order to recover such genes, we implemented a procedure in which we merge hits to the same gene occurring sequentially on the same genomic contig. If the distance between the start position of the first hit and the end position of the final hit gives ≥ 80% query coverage, the merge was considered valid.

Genes that were identified as single-copy (i.e., only one valid hit discovered from the entire genome) from ≥ 95% of the 1,587 species-representative assemblies were defined as candidate core genes, resulting in 62 genes.

### Profile HMM generation with AUGUSTUS

AUGUSTUS-PPX (36) provides a suite of scripts to generate block profile hidden Markov models (block profile HMMs; position-specific frequency matrices from a set of gap-less sequence blocks) from MSAs of homologous amino acid sequences, allowing sensitive and precise gene extraction from genome-scale data. We devised an iterative procedure using AUGUSTUS-PPX to build block profile HMMs with enriched homologous MSAs for 62 core gene candidates.

In each iteration, amino acid sequences of each gene are extracted from the species-representative assemblies with AUGUSTUS-PPX using the block profile HMMs from the previous iteration. Each extracted protein sequence was searched against the sequences from the respective MSA using MMseqs2. We accepted the protein sequence if its alignment covers at least 80% sequence length of a MSA member sequence. After each iteration a new MSA is generated by combining the previous and newly detected sequences using MAFFT v7.310 (37). The MSA is then used to build new block profile HMMs with AUGUSTUS-PPX.

For the first iteration, we used block profile HMMs built from the query centered MSAs (described above) for prediction and the amino acid sequences from Swiss-Prot with the corresponding gene annotation for homology search validation. We conducted three iterations of AUGUSTUS-PPX training for each of the core gene candidates we defined earlier, resulting in a final set of block profile HMMs.

### Quantifying the coverage of core gene candidates

To quantify the coverage of the core gene candidates on fungal species, we examined the presence of the genes from AUGUSTUS-PPX search against the species-representative genome assemblies. We repeated the final iteration of the profile HMM generation process described above to obtain the set of homologous protein sequences. The sequences with their alignment covering at least 80% sequence length of a member sequence of the respective MSA were accepted. A gene was defined present for the assemblies from which an accepted sequence was extracted.

For enhanced sensitivity, we relaxed the threshold and accepted the sequences covering at least 50% sequence length of a member. For the remaining sequences, the threshold was relaxed once more by accepting those aligned with E-value lower than 10^−3^.

We then tallied the proportion of the assemblies that reported the gene existing as a single-copy (i.e., only one homologous sequence detected), and those regardless of the copy number. Genes that ultimately failed to cover 95% of the species as a single-copy were rejected, while the remaining genes were defined as the final set of core marker genes.

To benchmark the core genes, we used the same method to quantify the existence coverage against 9,397 species-redundant genome assemblies, which passed the quality assessment but were unused because of their taxonomic redundancy.

### Canonical marker genes

We found the absence of the conventional marker genes for multi-gene phylogeny from our computational investigation, due to their functional divergence and lack of universality across the entire kingdom (38, 39). To supplement this, we collected a set of frequently used protein-coding phylogenetic markers from a review of fungal taxonomic literature, which we deemed canonical and included in the database. Profile HMM generation, coverage quantification, and benchmarking for these was performed identically as described for the core genes.

### Pipeline software development

We developed a bioinformatics pipeline integrating the process of marker gene extraction and phylogenetic analysis in a fully automated fashion. The modular pipeline allows users to process their biological sequences into sets of marker genes, align marker gene sequences, concatenate gene alignments, construct phylogenetic trees, and train their own marker MSAs and profile HMMs (Fig. 1b).

We developed a pipeline with three main modules: `profile`, `tree` and `train`. The `profile` module accepts genome, transcriptome and proteome data as input, and extracts marker gene sequences with AUGUSTUS-PPX using pre-trained block profile HMMs. In addition to the UFCG markers, we prepared profile HMMs for the ITS region and 758 single-copy orthologs from the fungal subset of OrthoDB v10 (24) available for the module. The module validates the sequences with a MMseqs2 search against the pre-defined homologous sequences, with stepwise relaxation of thresholds (coverage ≥ 80%, coverage ≥ 50%, E-value <10^−3^) as described above. A JSON file with valid amino acid and nucleotide sequences is produced as a result.

The `tree` module gathers the collection of JSON files produced with the `profile` module, constructs MSAs for each shared marker gene using MAFFT (37), removes alignment columns with a given gap threshold (default 50%), and generates phylogenetic trees in Newick format for the individual marker genes as well as from a concatenated MSA. For tree building, the user can choose among IQ-TREE (40), RAxML (41), and FastTree (42), with IQ-TREE being the default. Along with the bootstrap measure, the module computes a concatenation tree with branches annotated with Gene Support Index (GSI), the number of individual gene trees supporting the branch, as support values (21, 43).

Finally, the `train` module fully automates the iterative profile HMM generation process described above. The module accepts seed marker sequences and reference genome assemblies, and generates profile HMMs that can be directly utilised by the `profile` module.

### Phylogenetic tree construction

To test our database and pipeline, we utilised the UFCG marker genes to reconstruct the phylogenetic trees of fungal lineages (Fig. 3). Commands and parameters for the utilisation of our pipeline are described in Supplementary Table 2.

First, we downloaded 34 sequence datasets from the order Eurotiales, 13 genomic, 8 transcriptomic and 13 proteomic sequence sets (Supplementary Table 3) and extracted their UFCG marker genes with the `profile` module of our pipeline. Marker gene extraction was performed with UFCG v1.0 `profile` module, which utilises AUGUSTUS v3.4.0 and MMseqs2 v13.45111. The UFCG `tree` module automatically generated MSAs of the marker genes with MAFFT v7.310, removed alignment columns with ≥ 50% gaps, and drew the ML tree from the concatenation using JTT model (44) with IQ-TREE v2.0.3.

Additionally, we generated a kingdom-wide UFCG tree from genome assemblies of the entire 1,587 fungal species (Supplementary Fig. 2), with identical methods but using FastTree v2.1.10 (42) to generate the tree. Congruence of the major fungal lineages were compared against the kingdom-wide concatenation tree proposed by Li, et al. (45) and visualised as a tanglegram.

## Results and Discussion

### UFCG marker genes

We defined a set of 61 well-annotated and representative genes, namely UFCG marker genes (Supplementary Table 4). Determined by our computational pipeline, we included 41 core genes with 95% single-copy existence across 1,587 species-representative fungal genome assemblies. Additionally, we added the genes which have been frequently used to delineate higher-level classification of fungi (e.g., *RPB2, TEF1, TUB2* for phylum *Basidiomycota*) by fungal communities, resulting in 20 canonical genes (Supplementary Table 5).

Of the 62 candidate core genes, 41 covered more than 95% of the species-representative genomes in our dataset as single-copy (Fig. 2a). The remaining 21 failed the coverage threshold criterion and were rejected from the final set of core genes (Supplementary Fig. 1). When extended to the 9,397 species-redundant genome set, 40 of the 41 were identified as single-copy in more than 95% of the genomes (Fig. 2b).

**Fig. 2.**
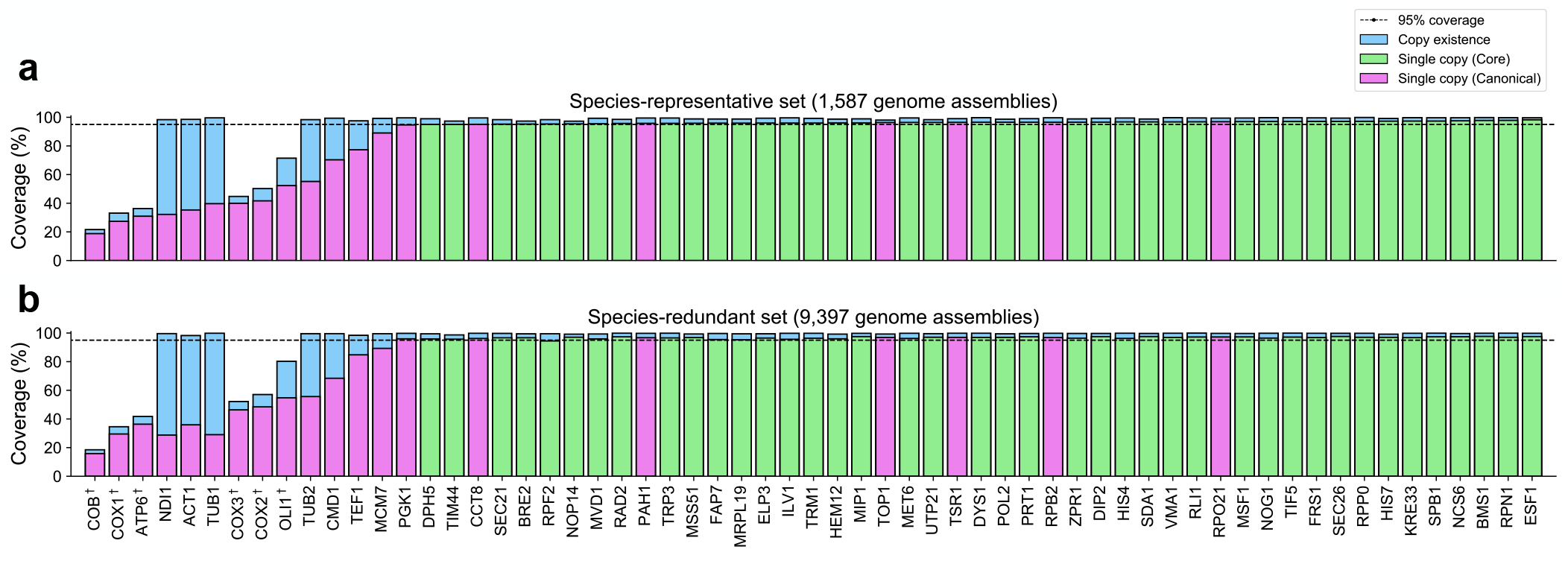
Existence coverage of 61 UFCG marker genes, represented as a proportion of fungal genome assemblies with a valid hit. (a) Coverage against 1,587 species-representative assemblies. (b) Coverage against 9,397 species-redundant assemblies. Presence of each marker gene against the given set of genome assemblies was identified using an AUGUSTUS-PPX search with their corresponding block profile HMMs. We then tallied the proportion of genome assemblies in which marker genes were i) present, regardless of copy-number (blue bars) and ii) present as single-copy (purple bars for canonical genes, green bars for core genes). Genes of mitochondrial origin (as annotated by the *Saccharomyces* genome database) were marked with a dagger (e.g., COX1^*†*^). Gene names are sorted by their single-copy coverage against the species-representative assemblies.

Meanwhile, only 7 out of 20 canonical genes met 95% single-copy threshold, while the main reason we speculate for this is the missing mitochondrial DNA in the genome assemblies. All but 6 canonical genes reported ≥ 98% coverage disregarding their copy numbers, in both the species-representative and the redundant set. These 6 genes are located on a mitochondrial genome, according to *Saccharomyces* genome database annotation (46), which most likely are universal genes that exists in more than 95% of species (47). However, 89.6% of the fungal genomes in the GenBank database are in a draft state (i.e., assembled below chromosome level) and therefore miss a certain fraction of DNA. We speculate that the mtDNA is especially affected by this since they might be deposited independently or might be rejected due to its uneven coverage in comparison to the remaining genomic DNA (48).

### Database contents

The UFCG gene database presents a summarised list of both core and canonical marker genes we defined, as well as descriptions of individual genes with downloadable resources (Fig. 1a, top). We prepared pre-trained block profile HMMs with both aligned and unaligned homologous amino acid sequences used to generate the models. Visualised MSAs are also available, constructed with the amino acid sequences extracted from 75 representative fungal species, which were implemented with MSAViewer (49). In addition, we offer direct links to the entries to external databases with corresponding annotations, including the *Saccharomyces* genome database (SGD; 46), UniProt (35) and NCBI Conserved Domain Database (CDD; 50).

The UFCG species database contains pre-extracted sequences of UFCG markers, ITS region, and BUSCO identified in the set of representative genome assemblies from 1,587 fungal species as described (Fig. 1a, bottom). Extracted sequences, metadata of originating genomes, and auxiliary run-time information were compiled into the JSON files (under field `data`, `genome_info`, `run_info`, respectively), which are downloadable from the database. We organised them into a sortable and searchable table, which provides the download links along with their NCBI accession numbers and taxonomic annotations. Additionally, we extracted UFCG markers, ITS region, and BUSCO sequences from 9,397 species-redundant genome assemblies, which are also downloadable from the database as compressed archives.

### Phylogenetic analysis with combined sequence types

The UFCG pipeline can extract marker genes from assorted types of biological sequences, including DNA, RNA, and protein. To illustrate this, we constructed a phylogenetic tree of UFCG marker genes extracted from 13 genomic, 8 transcriptomic, and 13 proteomic sequences, originating from three species under the order Eurotiales (Fig. 3). As shown by the topology of monophyletic clades grouped by their species origin, our gene database and pipeline successfully reconstructed the phylogenetic relationship from raw biological data in a fully automated procedure, regardless of their data types.

**Fig. 3.**
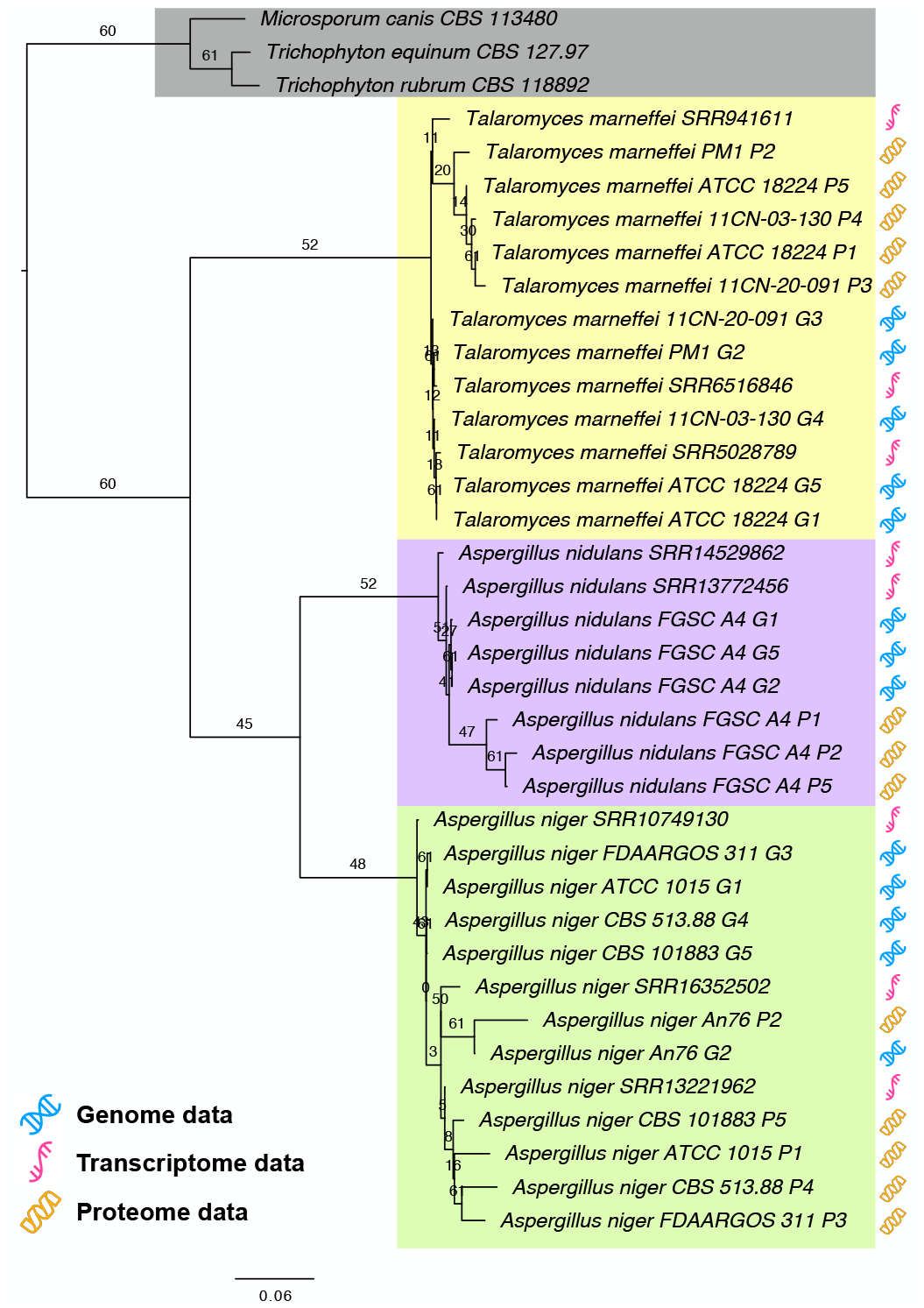
Maximum likelihood (ML) tree of the concatenated alignment of UFCG marker genes, extracted from either genomic, transcriptomic or proteomic data from 34 sequence datasets originated from three species under the order Eurotiales. As outgroup we included three species from the order Onygenales (highlighted in grey). Branches of the resulting tree were annotated by their GSI values. Monophyletic clades clustered by their species origin were highlighted with coloured box (Yellow, *Talaromyces marneffei* ; Purple, *Aspergillus nidulans*; Green, *Aspergillus niger*). Type of sequence origin was marked with the respective symbol (refer to the legend).

### Kingdom-wide phylogenetic reconstruction

One of the key advantages of the UFCG pipeline is its ability to automatically construct phylogenies from computationally detectable core genes. This is particularly useful for larger genome datasets, where manual extraction of marker genes is prohibitive.

We reconstructed a kingdom-wide phylogenetic relationship of 1,587 species-representative assemblies using UFCG marker genes extracted with our pipeline (Supplementary Fig. 2). From the tanglegram comparing the topology between our tree and proposed genome-scale fungal phylogeny using BUSCO sequences (45), 14 out of 18 major lineages of fungi were congruently placed (Supplementary Fig. 3). Although two incongruent pairs were observed (*Wallemiomycotina* and *Ustilagomycotina*; *Glomeromycotina* and *Mortierellomycotina*), we were able to find other studies supporting our phylogeny with congruent placement of pairs (51, 52, respectively).

## Concluding remarks

Our novel database of fungal marker genes and pipeline provides a robust and easy-to-use method for genome-wide phylogenetic analysis of fungi. Similar approaches for prokaryotic communities such as Genome Taxonomy Database (GTDB; 19), AutoMLST (20), and UBCG (21) have shown the value that automatic phylogenetic analysis brings. As the first fungal core gene database with an automated phylogenetic pipeline, we expect UFCG to be of similar interest and help to tackle the challenge of genome-scale fungal phylogenetic analysis.

## Data Availability

The UFCG database is freely available without registration at https://ufcg.steineggerlab.com. Entire content of the database is licensed under CC BY-SA 4.0. The pipeline is implemented in Java and is available as GPLv3 licensed free opensource software at https://github.com/endixk/ufcg.

## Acknowledgements

We thank the members of Seoul National University Fungal EcoPhylogeny Laboratory, especially Young Woon Lim, Changwan Seo, Ki Hyeong Park, and Myung Soo Park for discussions and advice. We thank Seong-In Na for his kind permission to include the source code he developed as well as Milot Mirdita for helping to revise the manuscript. M.S. acknowledges support from the National Research Foundation of Korea (NRF), grants [2019R1A6A1A10073437, 2020M3A9G7103933, 2021R1C1C102065, 2021M3A9I4021220], and the Creative-Pioneering Researchers Program through Seoul National University.

## Conflict of interest statement

None declared.

## Notes

https://ufcg.steineggerlab.com

### Competing Interest Statement

The authors have declared no competing interest.

