## Supplementary materials for "UFCG: database of universal fungal core genes and pipeline for genome-wide phylogenetic analysis of fungi"

### Tables

- Supplementary Table 1. Statistics of AUGUSTUS gene prediction of species models from 5 fungal phyla.
- Supplementary Table 2. Commands and parameters for UFCG pipeline to generate the UFCG trees.
- Supplementary Table 3. Description of 34 sequences originated from 3 species under the order Eurotiales.
- Supplementary Table 4. Detailed information of 61 UFCG marker genes.
- Supplementary Table 5. List of 20 canonical marker genes.

### Figures

- Supplementary Figure 1. Existence coverage of 21 candidate core marker genes below 95% single copy proportion.
- Supplementary Figure 2. Topology of the maximum likelihood tree of 1,587 genome asseblies representing fungal species.
- Supplementary Figure 3. Tanglegram comparing the topologies of two kingdom-wide tree of fungal species.

**Supplementary Table 1.** Statistics of AUGUSTUS gene prediction count from 1,587 species-representative genome assemblies, using pre-trained species models from 5 fungal phyla.

| Species | Phylum | Average prediction count (±SE) |
|---|---|---|
| *Encephalitozoon cuniculi* | Rozellomycota | 4128.53 (±76.2312) |
| *Conidiobolus coronatus* | Entomophthoromycota | 11478.4 (±194.335) |
| *Rhizopus oryzae* | Mucoromycota | **13550.8** (±215.982) |
| *Saccharomyces cerevisiae* | Ascomycota | 7625.03 (±99.7421) |
| *Cryptococcus neoformans* | Basidiomycota | 9009.65 (±123.441) |

**Supplementary Table 2.** Commands and parameters for UFCG pipeline to generate the UFCG trees from 34 Eurotiales species and 1,587 fungal assemblies

| Task | Command |
|---|---|
| Marker gene extraction from genomes | `java -jar UFCG.jar profile -i [FASTA] -o [OUT] --info [METADATA]` |
| Marker gene extraction from transcriptomes | `java -jar UFCG.jar profile-rna -p 1 -i [FASTQ] -o [OUT] --info [METADATA]` |
| Marker gene extraction from proteomes | `java -jar UFCG.jar profile-pro -i [FASTA] -o [OUT] --info [METADATA]` |
| Eurotiales species tree generation | `java -jar UFCG.jar tree -i [IN] -l label -o [OUT] -a protein -p iqtree` |
| Kingdom-wide tree generation | `java -jar UFCG.jar tree -i [IN] -l label -o [OUT] -a protein -p fasttree` |

**Supplementary Table 3.** Description of 34 genomic, transcriptomic, and proteomic sequences retrieved from NCBI, originated from 3 species under the order Eurotiales: *Talaromyces marneffei*, *Aspergillus nidulans*, and *Aspergillus niger*.

| Species | Sequence type | Label | NCBI accession |
|---|---|---|---|
| *Talaromyces marneffei* | Genome | Talaromyces marneffei ATCC 18224 G1 | GCA_000001985.1 |
| *Talaromyces marneffei* | Genome | Talaromyces marneffei PM1 G2 | GCA_000750115.1 |
| *Talaromyces marneffei* | Genome | Talaromyces marneffei 11CN-20-091 G3 | GCA_009556855.1 |
| *Talaromyces marneffei* | Genome | Talaromyces marneffei 11CN-03-130 G4 | GCA_009650675.1 |
| *Talaromyces marneffei* | Genome | Talaromyces marneffei ATCC 18224 G5 | GCF_000001985.1 |
| *Talaromyces marneffei* | Proteome | Talaromyces marneffei ATCC 18224 P1 | GCA_000001985.1 |
| *Talaromyces marneffei* | Proteome | Talaromyces marneffei PM1 P2 | GCA_000750115.1 |
| *Talaromyces marneffei* | Proteome | Talaromyces marneffei 11CN-20-091 P3 | GCA_009556855.1 |
| *Talaromyces marneffei* | Proteome | Talaromyces marneffei 11CN-03-130 P4 | GCA_009650675.1 |
| *Talaromyces marneffei* | Proteome | Talaromyces marneffei ATCC 18224 P5 | GCF_000001985.1 |
| *Talaromyces marneffei* | Transcriptome | Talaromyces marneffei SRR5028789 | SRR5028789 |
| *Talaromyces marneffei* | Transcriptome | Talaromyces marneffei SRR6516846 | SRR6516846 |
| *Talaromyces marneffei* | Transcriptome | Talaromyces marneffei SRR941611 | SRR941611 |
| *Aspergillus nidulans* | Genome | Aspergillus nidulans FGSC A4 G1 | GCA_000011425.1 |
| *Aspergillus nidulans* | Genome | Aspergillus nidulans FGSC A4 G2 | GCA_000149205.2 |
| *Aspergillus nidulans* | Genome | Aspergillus nidulans FGSC A4 G5 | GCF_000149205.2 |
| *Aspergillus nidulans* | Proteome | Aspergillus nidulans FGSC A4 P1 | GCA_000011425.1 |
| *Aspergillus nidulans* | Proteome | Aspergillus nidulans FGSC A4 P2 | GCA_000149205.2 |
| *Aspergillus nidulans* | Proteome | Aspergillus nidulans FGSC A4 P5 | GCF_000149205.2 |
| *Aspergillus nidulans* | Transcriptome | Aspergillus nidulans SRR13772456 | SRR13772456 |
| *Aspergillus nidulans* | Transcriptome | Aspergillus nidulans SRR14529862 | SRR14529862 |
| *Aspergillus niger* | Genome | Aspergillus niger ATCC 1015 G1 | GCA_000230395.2 |
| *Aspergillus niger* | Genome | Aspergillus niger An76 G2 | GCA_001515345.1 |
| *Aspergillus niger* | Genome | Aspergillus niger FDAARGOS_311 G3 | GCA_002211485.2 |
| *Aspergillus niger* | Genome | Aspergillus niger CBS 513.88 G4 | GCF_000002855.3 |
| *Aspergillus niger* | Genome | Aspergillus niger CBS 101883 G5 | GCF_003184595.1 |
| *Aspergillus niger* | Proteome | Aspergillus niger ATCC 1015 P1 | GCA_000230395.2 |
| *Aspergillus niger* | Proteome | Aspergillus niger An76 P2 | GCA_001515345.1 |
| *Aspergillus niger* | Proteome | Aspergillus niger FDAARGOS_311 P3 | GCA_002211485.2 |
| *Aspergillus niger* | Proteome | Aspergillus niger CBS 513.88 P4 | GCF_000002855.3 |
| *Aspergillus niger* | Proteome | Aspergillus niger CBS 101883 P5 | GCF_003184595.1 |
| *Aspergillus niger* | Transcriptome | Aspergillus niger SRR10749130 | SRR10749130 |
| *Aspergillus niger* | Transcriptome | Aspergillus niger SRR13221962 | SRR13221962 |
| *Aspergillus niger* | Transcriptome | Aspergillus niger SRR16352502 | SRR16352502 |

**Supplementary Table 4.** Detailed information of 61 UFCG marker genes. Genes are labeled by their *Saccharomyces* genome database (SGD) names.

| Gene | Type | SGD ID | UniProt ID | CDD ID | COG* | Function |
|---|---|---|---|---|---|---|
| *ACT1* | Canonical | YFL039C | P60010 | KOG0676 | Z | $\gamma$-Actin |
| *ATP6* | Canonical | Q0085 | P00854 | KOG4665 | C | $F_1F_0$ ATP synthase subunit 6 |
| *BMS1* | Core | YPL217C | Q08965 | KOG1951 | J | Ribosome biogenesis protein |
| *BRE2* | Core | YLR015W | P43132 | KOG2626 | B/K | COMPASS component |
| *CCT8* | Canonical | YJL008C | P47079 | KOG0362 | O | Chaperonin-containing T-complex subunit $\theta$ |
| *CMD1* | Canonical | YBR109C | P06787 | KOG0027 | T | Calmodulin |
| *COB* | Canonical | Q0105 | P00163 | KOG4663 | C | Cytochrome b |
| *COX1* | Canonical | Q0045 | P00401 | KOG4769 | C | Cytochrome c oxidase subunit 1 |
| *COX2* | Canonical | Q0250 | P00410 | KOG4767 | C | Cytochrome c oxidase subunit 2 |
| *COX3* | Canonical | Q0275 | P00420 | KOG4664 | C | Cytochrome c oxidase subunit 3 |
| *DIP2* | Core | YLR129W | Q12220 | KOG0306 | A | U3 small nucleolar RNA-associated protein 12 |
| *DPH5* | Core | YLR172C | P32469 | KOG3123 | J | Diphthine methyl ester synthase |
| *DYS1* | Core | YHR068W | P38791 | KOG2924 | O | Deoxyhypusine synthase |
| *ELP3* | Core | YPL086C | Q02908 | KOG2535 | B/K | Elongator complex protein 3 |
| *ESF1* | Core | YDR365C | Q06344 | KOG2318 | S | Pre-rRNA-processing protein |
| *FAP7* | Core | YDL166C | Q12055 | KOG3347 | F | Adenylate kinase isoenzyme 6 homolog |
| *FRS1* | Core | YLR060W | P15624 | KOG2472 | J | Phenylalanine–tRNA ligase beta subunit |
| *HEM12* | Core | YDR047W | P32347 | KOG2872 | H | Uroporphyrinogen decarboxylase |
| *HIS4* | Core | YCL030C | P00815 | KOG2697 | E | Histidine biosynthesis trifunctional protein |
| *HIS7* | Core | YBR248C | P33734 | KOG0623 | E | Imidazole glycerol phosphate synthase |
| *ILV1* | Core | YER086W | P00927 | KOG1250 | E | Threonine dehydratase |
| *KRE33* | Core | YNL132W | P53914 | KOG2036 | R | RNA cytidine acetyltransferase |
| *MCM7* | Canonical | YBR202W | P38132 | KOG0482 | L | Mini-chromosome maintenance complex subunit |
| *MET6* | Core | YER091C | P05694 | KOG2263 | E | 5-methyltetrahydropteroyltriglutamate–homocysteine methyltransferase |
| *MIP1* | Core | YOR330C | P15801 | KOG3657 | L | DNA polymerase $\gamma$ |
| *MRPL19* | Core | YNL185C | P53875 | KOG3257 | J | 54S ribosomal protein L19 |
| *MSF1* | Core | YPR047W | P08425 | KOG2783 | J | Phenylalanine–tRNA ligase |
| *MSS51* | Core | YLR203C | P32335 | - | O | Mitochondrial splicing suppressor protein 51 |
| *MVD1* | Core | YNR043W | P32377 | KOG2833 | I | Diphosphomevalonate decarboxylase |
| *NCS6* | Core | YGL211W | P53088 | KOG2840 | R | Cytoplasmic tRNA 2-thiolation protein 1 |
| *NDI1* | Canonical | YML120C | P32340 | KOG2495 | C | NADH-ubiquinone reductase |
| *NOG1* | Core | YPL093W | Q02892 | KOG1490 | R | Nucleolar GTP-binding protein 1 |
| *NOP14* | Core | YDL148C | Q99207 | KOG2147 | J | Nucleolar complex protein 14 |
| *OLI1* | Canonical | Q0130 | P61829 | KOG3025 | C | $F_1F_0$ ATP synthase subunit 9 |
| *PAH1* | Canonical | YMR165C | P32567 | KOG2116 | N/I | Phosphatidate phosphatase |
| *PGK1* | Canonical | YCR012W | P00560 | KOG1367 | G | Phosphoglycerate kinase |
| *POL2* | Core | YNL262W | P21951 | KOG1798 | L | DNA polymerase epsilon catalytic subunit A |
| *PRT1* | Core | YOR361C | P06103 | KOG2314 | J | Eukaryotic translation initiation factor 3 subunit B |
| *RAD2* | Core | YGR258C | P07276 | KOG2520 | L | DNA repair protein |
| *RLI1* | Core | YDR091C | Q03195 | KOG0063 | A | Translation initiation factor |
| *RPB2* | Canonical | YOR151C | P08518 | KOG0214 | K | DNA-directed RNA polymerase II core subunit |
| *RPF2* | Core | YKR081C | P36160 | KOG3031 | J | Ribosome biogenesis protein |
| *RPN1* | Core | YHR027C | P38764 | KOG2005 | O | 26S proteasome regulatory subunit |
| *RPO21* | Canonical | YDL140C | P04050 | KOG0260 | K | DNA-directed RNA polymerase II core subunit |
| *RPP0* | Core | YLR340W | P05317 | KOG0815 | J | 60S acidic ribosomal protein P0 |
| *SDA1* | Core | YGR245C | P53313 | KOG2229 | D/Z | Severe depolymerization of actin protein 1 |
| *SEC21* | Core | YNL287W | P32074 | KOG1078 | U | Coatomer subunit gamma |
| *SEC26* | Core | YDR238C | P41810 | KOG1058 | U | Coatomer subunit beta |
| *SPB1* | Core | YCL054W | P25582 | KOG1098 | A/R | 27S pre-rRNA (guanosine$_{2922}$-2'-O)-methyltransferase |
| *TEF1* | Canonical | YPR080W | P02994 | KOG0052 | J | Translation elongation factor EF-1$\alpha$ |
| *TIF5* | Core | YPR041W | P38431 | KOG2767 | J | Eukaryotic translation initiation factor 5 |
| *TIM44* | Core | YIL022W | Q01852 | KOG2580 | U | Mitochondrial import inner membrane translocase subunit |
| *TOP1* | Canonical | YOL006C | P04786 | KOG0981 | L | DNA topoisomerase 1 |
| *TRM1* | Core | YDR120C | P15565 | KOG1253 | J | tRNA (guanine$_{26}$-$N_2$)-dimethyltransferase |
| *TRP3* | Core | YKL211C | P00937 | KOG0026 | E | Multifunctional tryptophan biosynthesis protein |
| *TSR1* | Canonical | YDL060W | Q07381 | KOG1980 | S | Ribosome maturation factor |
| *TUB1* | Canonical | YML085C | P09733 | KOG1376 | Z | $\alpha$-tubulin |
| *TUB2* | Canonical | YFL037W | P02557 | KOG1375 | Z | $\beta$-tubulin |
| *UTP21* | Core | YLR409C | Q06078 | KOG1539 | R | U3 small nucleolar RNA-associated protein 21 |
| *VMA1* | Core | YDL185W | P17255 | KOG1540 | H | V-type proton ATPase catalytic subunit A |
| *ZPR1* | Core | YGR211W | P53303 | KOG2703 | R | Zinc finger protein |

*COG, clusters of orthologous group: A, RNA processing and modification; B, Chromatin Structure and dynamics; C, Energy production and conversion; D, Cell cycle control and mitosis; E, Amino Acid metabolis and transport; F, Nucleotide metabolism and transport; G, Carbohydrate metabolism and transport; H, Coenzyme metabolis; I, Lipid metabolism; J, Tranlsation; K, Transcription; L, Replication and repair; N, Cell motility; O, Post-translational modification, protein turnover, chaperone functions; T, Signal Transduction; U, Intracellular trafficing and secretion; Z, Cytoskeleton; R, General Functional Prediction only; S, Function Unknown.

**Supplementary Table 5.** List of 20 canonical marker genes with example fungal taxa with phylogenetic analysis using the markers (see also 1; 2).

| Gene | Aliases | Example taxa | References |
|---|---|---|---|
| *ACT1* | *ACT* | *Cryptococcus*, *Glomeromycota* | 3, 4 |
| *ATP6* | - | *Boteales*, *Agaricus* | 5, 6 |
| *CCT8* | *TCP1θ* | *Aspergillus*, *Saccharomyces* | 7, 8 |
| *CMD1* | *CAL, CaM* | *Eurotiales*, *Penicillium* | 9, 10 |
| *COB* | - | *Aspergillus*, *Glomeromycota* | 11, 12 |
| *COX1* | - | *Pezizomycotina*, *Glomeromycota* | 12, 13 |
| *COX2* | - | *Peronosporomycetes* | 14 |
| *COX3* | - | *Boteales* | 5 |
| *MCM7* | *CDC47* | *Ascomycota*, *Kickxellomycotina* | 15, 16 |
| *NDI1* | *NAD1-6* | *Beauveria*, *Glomeromycota* | 12, 17 |
| *OLI1* | *mtATP9* | *Beauveria*, *Glomeromycota* | 12, 17 |
| *PAH1* | *LNS2* | *Pucciniomycota* | 1 |
| *PGK1* | *PGK* | *Fusarium*, *Penicillium* | 1, 18 |
| *RPB2* | - | *Ascomycota*, *Basidiomycota* | 19, 20 |
| *RPO21* | *RPB1* | *Inocybe*, *Zygomycota* | 21, 22 |
| *TEF1* | *TEF1α* | *Basidiomycota*, *Zygomycota* | 19, 22 |
| *TOP1* | - | *Fusarium*, *Penicillium* | 1, 23 |
| *TSR1* | - | *Kickxellomycotina* | 15 |
| *TUB1* | - | *Microsporidia* | 24 |
| *TUB2* | *BenA* | *Basidiomycota*, *Microsporidia* | 24, 25 |

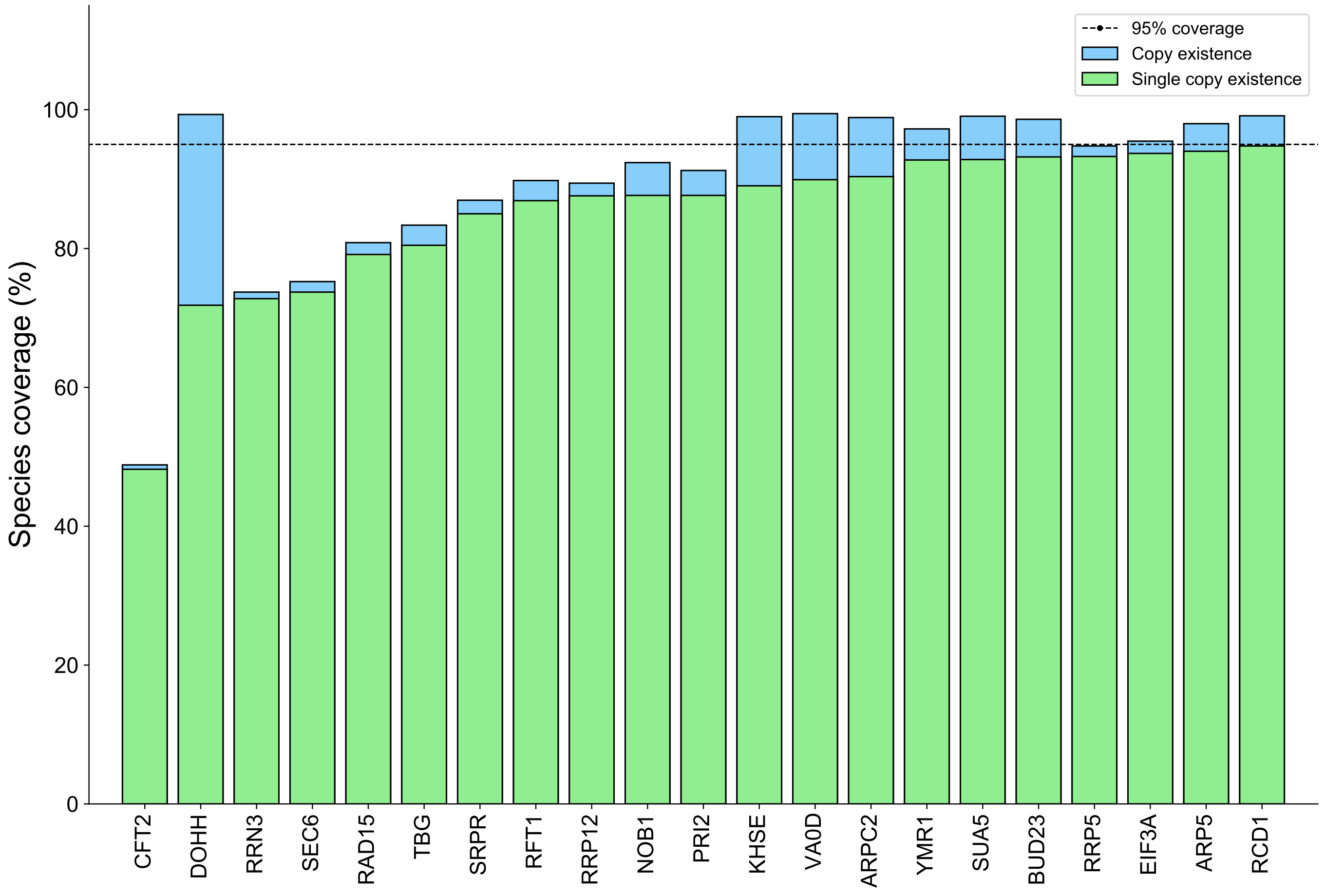


**Supplementary Figure 1.** Existence coverage of 21 candidate core marker genes, which failed to achieve 95% single copy proportion of covered entries among the 1,587 genome assemblies representing fungal species.

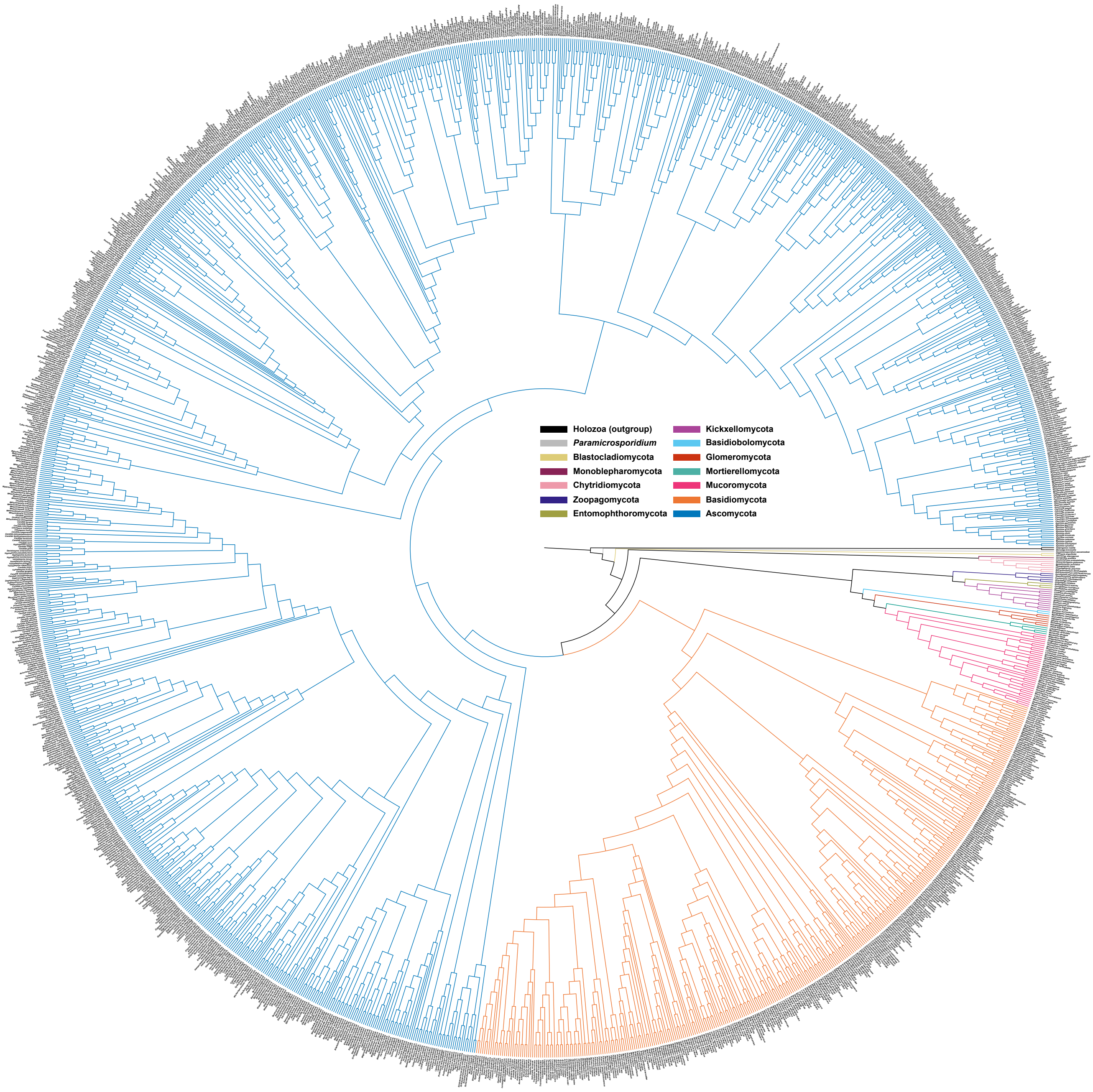


**Supplementary Figure 2.** Topology of the maximum likelihood tree of 1,587 genome assemblies representing fungal species. Tree was generated from the concatenated amino acid sequence alignment of 61 UFCG marker genes, using FastTree v2.1.10. Branches are coloured based on their phylum (refer to the legend).

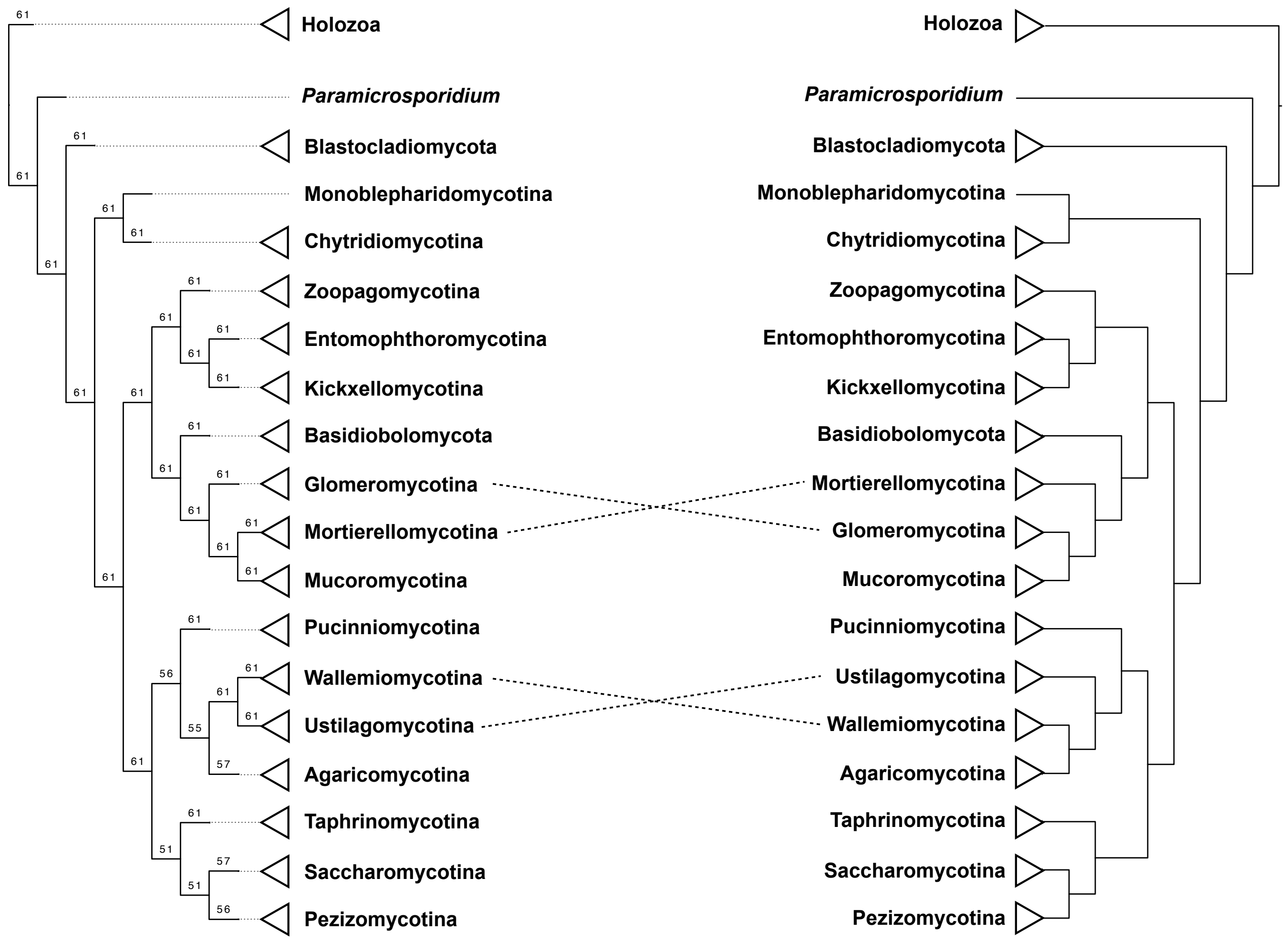


**Supplementary Figure 3.** Tanglegram comparing the topologies of two kingdom-wide tree of fungal species: Left, UFCG marker gene concatenation tree; Right, BUSCO concatenation tree presented by Li, et al. (26). Branches of UFCG trees were annotated by their gene support index (GSI) values. Discrepancies between the trees were visualised by dotted lines connecting the corresponding clades.
